# An improved reversibly dimerizing mutant of the FK506-binding protein FKBP

**DOI:** 10.1101/053751

**Authors:** Juan J. Barrero, Effrosyni Papanikou, Jason C. Casler, Kasey J. Day, Benjamin S. Glick

**Author notes:** Contributed equally.

## Abstract

FK506-binding protein (FKBP) is a monomer that binds to FK506, rapamycin, and related ligands. The F36M substitution, in which Phe36 in the ligand-binding pocket is changed to Met, leads to formation of antiparallel FKBP dimers, which can be dissociated into monomers by ligand binding. This FKBP(M) mutant has been employed in the mammalian secretory pathway to generate aggregates that can be dissolved by ligand addition to create cargo waves. However, when testing this approach in yeast, we found that dissolution of FKBP(M) aggregates was inefficient. An improved reversibly dimerizing FKBP formed aggregates that dissolved more readily. This FKBP(L,V) mutant carries the F36L mutation, which increases the affinity of ligand binding, and the I90V mutation, which accelerates ligand-induced dissociation of the dimers. The FKBP(L,V) mutant expands the utility of reversibly dimerizing FKBP.

## Introduction

Human FKBP, also known as FKBP12, is a monomeric 12-kDa *cis/trans* peptidyl-prolyl isomerase and a member of a family of proteins that bind the immunosuppressive drugs FK506 and rapamycin (Galat, 2013). The FKBP-FK506 complex associates with the phosphatase calcineurin, whereas the FKBP-rapamycin complex associates with the FKBP-rapamycin binding (FRB) domain of the kinase mTOR. As a research tool, the FKBP-rapamycin-FRB interaction has been widely used for ligand-inducible heterodimerization of fusion proteins (Fegan *et al.*, 2010). The cytotoxicity of rapamycin can be avoided by using synthetic ligands that bind to wild-type or mutant forms of FKBP.

When Phe36 in the ligand-binding pocket of FKBP was changed to Met to create the FKBP(M) mutant, a novel property emerged (Rollins *et al.*, 2000). The altered ligand-binding interface self-associated so that FKBP(M) formed an antiparallel homodimer, and in the presence of ligand, dimerization was blocked. Expression in mammalian cells of enhanced GFP (EGFP) fused to four tandem copies of FKBP(M) resulted in formation of fluorescent aggregates, presumably due to a heterogeneous set of cross-associations between the EGFP-FKBP(M)x4 molecules. Those aggregates could be dissolved by adding ligand.

Reversible aggregation of FKBP(M) has been exploited to create fluorescent cargo waves in the mammalian secretory pathway (Rivera *et al.*, 2000). When EGFP-FKBP(M)x4 was targeted to the lumen of the endoplasmic reticulum (ER), it formed aggregates that were too large to exit the ER. Addition of ligand dissolved the aggregates to yield monomers that were incorporated into ER-derived vesicles and transported to the Golgi. Variations on this method have been used to test models for protein traffic through the Golgi stack (Volchuk *et al.*, 2000; Lavieu *et al.*, 2013; Rizzo *et al.*, 2013).

We are interested in using reversible aggregation to create cargo waves in the secretory pathway of the yeast *Saccharomyces cerevisiae*, which is a popular model organism for studying membrane traffic (Kaiser *et al.*, 1997; Papanikou and Glick, 2009). No good method is yet available for tracking the movement of fluorescent cargoes through the yeast secretory pathway. We observed that when fluorescent proteins were targeted to the yeast ER, GFP variants exited the ER slowly (Fitzgerald and Glick, 2014), but the highly soluble tetrameric DsRed variant E2-Crimson (Strack *et al.*, 2009) exited the ER much faster (E. Papanikou, unpublished observations). E2-Crimson was therefore deemed a good starting point for engineering a fluorescent secretory cargo. Fusion of a single copy of FKBP(M) to tetrameric E2-Crimson was predicted to generate fluorescent aggregates that could be dissolved by adding ligand (Figure 1).

**FIGURE 1:**
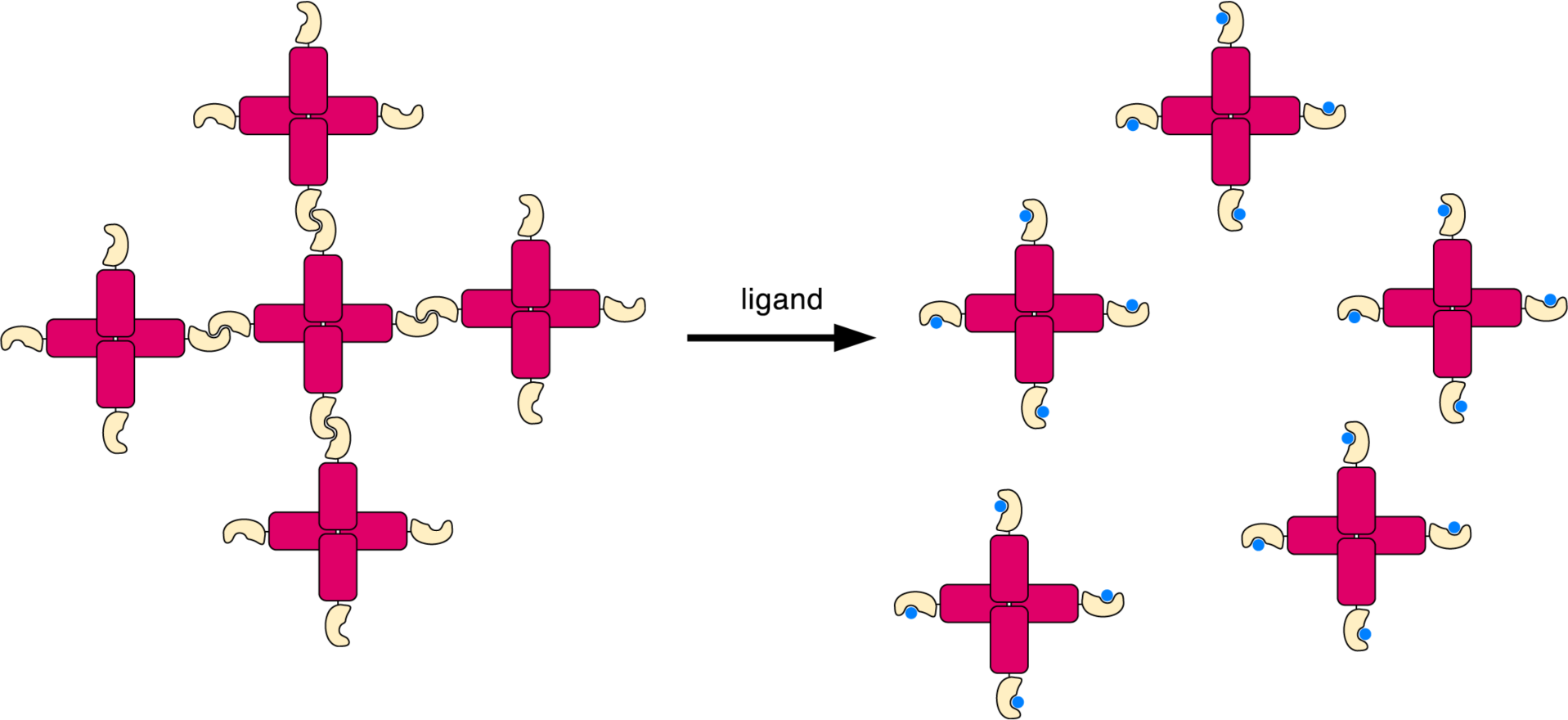
Strategy for generating fluorescent aggregates using dimerizing FKBP. Tetrameric E2-Crimson (red) is fused to a dimerizing FKBP mutant (gold) to generate cross-linked aggregates. Addition of ligand (blue) disrupts FKBP dimerization, thereby yielding soluble tetramers.

For initial characterization, we fused FKBP(M) to the C-terminus of E2-Crimson and expressed this construct in the yeast cytosol. Fluorescent aggregates formed and could be dissolved by adding ligand. However, dissolution of the aggregates was relatively slow and required high ligand concentrations. To improve this system, we engineered an alternative dimerizing FKBP mutant that permits faster dissolution of aggregates with lower concentrations of ligand.

## Results and Discussion

### Generation of fluorescent aggregates in the yeast cytosol

It was previously shown that expression in mammalian cells of EGFP fused to four tandem copies of FKBP(M) resulted in formation of fluorescent aggregates (Rollins *et al.*, 2000). To replicate this effect in yeast, we expressed EGFP-FKBP(M)x4 from an integrating vector using the strong constitutive *TPI1* promoter. Many of the cells showed one or two fluorescent aggregates (Figure 2A).

**FIGURE 2:**
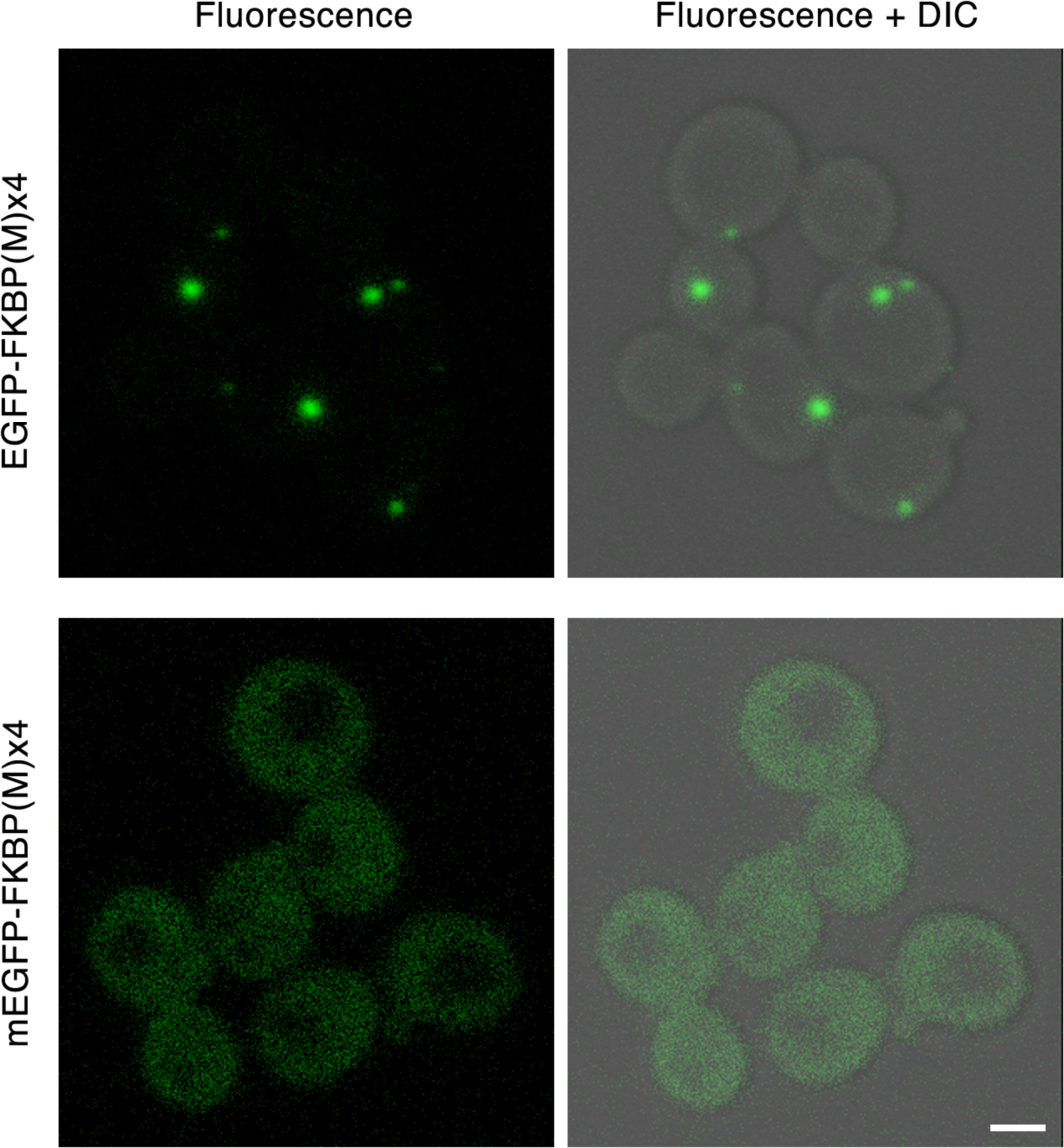
Influence of the fluorescent protein tag on the formation of aggregates containing four tandem copies of FKBP(M). Yeast cells were transformed with integrating vectors to express either EGFP-FKBP(M)x4 or mEGFP-FKBP(M)x4. Green fluorescence and differential interference contrast (DIC) images of logarithmically growing cells were captured by confocal microscopy, using the same parameters for both strains. Representative flourescence and merged images are shown. To enhance weaker signals, the gamma value of the fluorescence images was adjusted to 2.0 using Adobe Photoshop. Scale bar, 2 μm.

EGFP itself has a weak tendency to dimerize, and this property can dramatically affect the localization of certain fusion proteins (Zacharias *et al.*, 2002; Lisenbee *et al.*, 2003; Snapp *et al.*, 2003). We therefore made an alternative yeast expression construct in which EGFP was replaced with mEGFP, which contains the monomerizing A206K mutation (Zacharias *et al.*, 2002). Yeast cells expressing mEGFP-FKBP(M)x4 showed a diffuse cytosolic fluorescence with few if any aggregates (Figure 2B). We infer that aggregate formation with EGFP-FKBP(M)x4 likely involves EGFP dimerization as well as FKBP(M) dimerization.

A conceptually simpler way to generate fluorescent aggregates is to fuse a single copy of FKBP(M) to a tetrameric fluorescent protein (see Figure 1). When FKBP(M) was fused to the C-terminus of E2-Crimson and expressed in yeast, many of the cells showed one or two fluorescent aggregates of varying size (Figure 3, top panels). By contrast, when FKBP(M) was replaced with wild-type FKBP, none of the cells had fluorescent aggregates (data not shown). Thus, the E2-Crimson-FKBP(M) construct was deemed promising for the generation of reversible aggregates in yeast cells.

**FIGURE 3:**
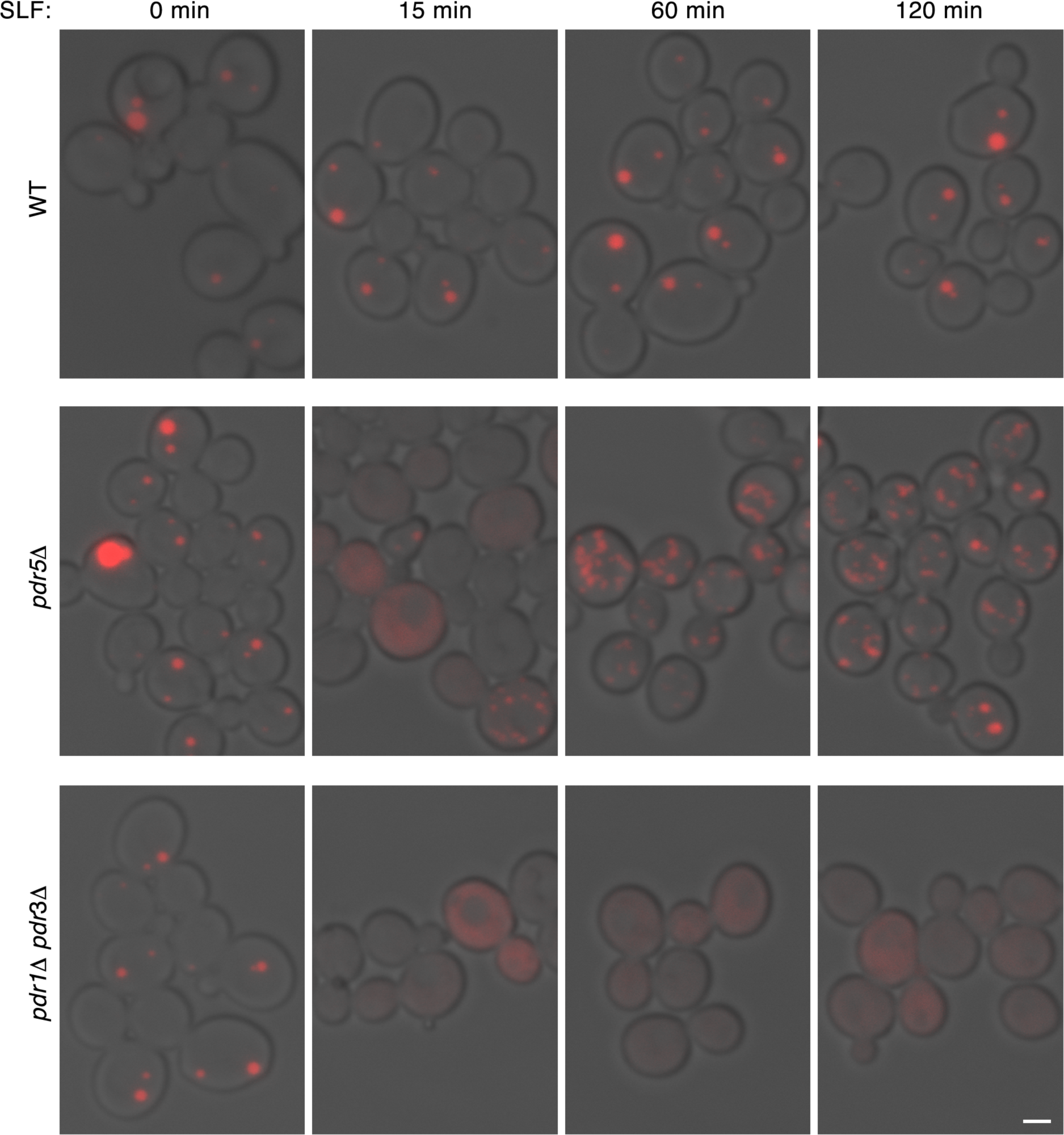
Enhanced aggregate dissolution in drug-sensitive yeast strains. An integrating vector encoding E2-Crimson-FKBP(M) was transformed into isogenic wild-type and *pdr5Δ* and *pdr1Δ pdr3Δ* strains. Logarithmically growing cultures were treated with 250 μm SLF for 0 to 120 min as indicated, and were imaged by confocal microscopy to capture red fluorescence and DIC images, using the same parameters in all cases. Representative merged images are shown. Scale bar, 2 μm.

### Dissolution of aggregates in a drug-sensitive yeast strain

In an effort to dissolve the E2-Crimson-FKBP(M) aggregates, we used a commercially available synthetic ligand of FKBP called SLF (Holt *et al.*, 1993). However, when wild-type yeast cells expressing E2-Crimson-FKBP(M) were incubated with 250 μM SLF for up to 2 h, no significant dissolution of the aggregates was seen (Figure 3, top panels).

We reasoned that the cells might be removing SLF with the aid of ABC-type pleiotropic drug transporter (PDR) proteins (Rogers *et al.*, 2001). One of the major drug transporters in *S. cerevisiae* is Pdr5. In a *pdr5Δ* mutant expressing E2-Crimson-FKBP(M), treatment for 15 min with SLF led to substantial dissolution of the aggregates, with many cells showing a uniform weak cytosolic fluorescence and others showing cytosolic fluorescence plus very small aggregates (Figure 3, middle panels). But after 1 h the cytosolic fluorescence was no longer visible, and the cells contained multiple small aggregates. Evidently, the cells adapted over time (Schüller *et al.*, 2007; Thakur *et al.*, 2008) by increasing their capacity to export SLF using transporters other than Pdr5.

Expression of Pdr5 and many other PDR proteins in *S. cerevisiae* is controlled by the Pdr1 and Pdr3 pair of transcription factors (Rogers *et al.*, 2001). To achieve a broad and persistent suppression of drug export, we disrupted both the *PDR1* and *PDR3* genes. In a *pdr1Δ pdr3Δ* double mutant expressing E2-Crimson-FKBP(M), addition of SLF dissolved the aggregates to generate a uniform cytosolic fluorescence that persisted for at least 2 h (Figure 3, bottom panels, and data not shown). The *pdr1Δ pdr3Δ* strain was therefore chosen as the starting point for further analysis.

### Enhancement of the ligand effect with the F36L mutation

To dissolve E2-Crimson-FKBP(M) aggregates, we used SLF at 250 μM because lower concentrations were less effective. A *pdr1Δ pdr3Δ* strain showed no growth defect in medium containing 250 μM SLF (data not shown), but the use of such high drug concentrations is expensive and raises concerns about nonspecific effects on the cells. These issues could be addressed by using a dimerizing FKBP variant with stronger ligand binding.

The F36M mutation reduces ligand binding affinities approximately 10-fold (Rollins *et al.*, 2000), whereas the F36L mutation has only a minimal effect on ligand binding (DeCenzo *et al.*, 1996). We therefore tested whether dimerization occurred with an F36L mutant of FKBP, here termed FKBP(L). Indeed, E2-Crimson-FKBP(L) formed cytosolic aggregates similar to those seen with E2-Crimson-FKBP(M) (Figure 4).

**FIGURE 4:**
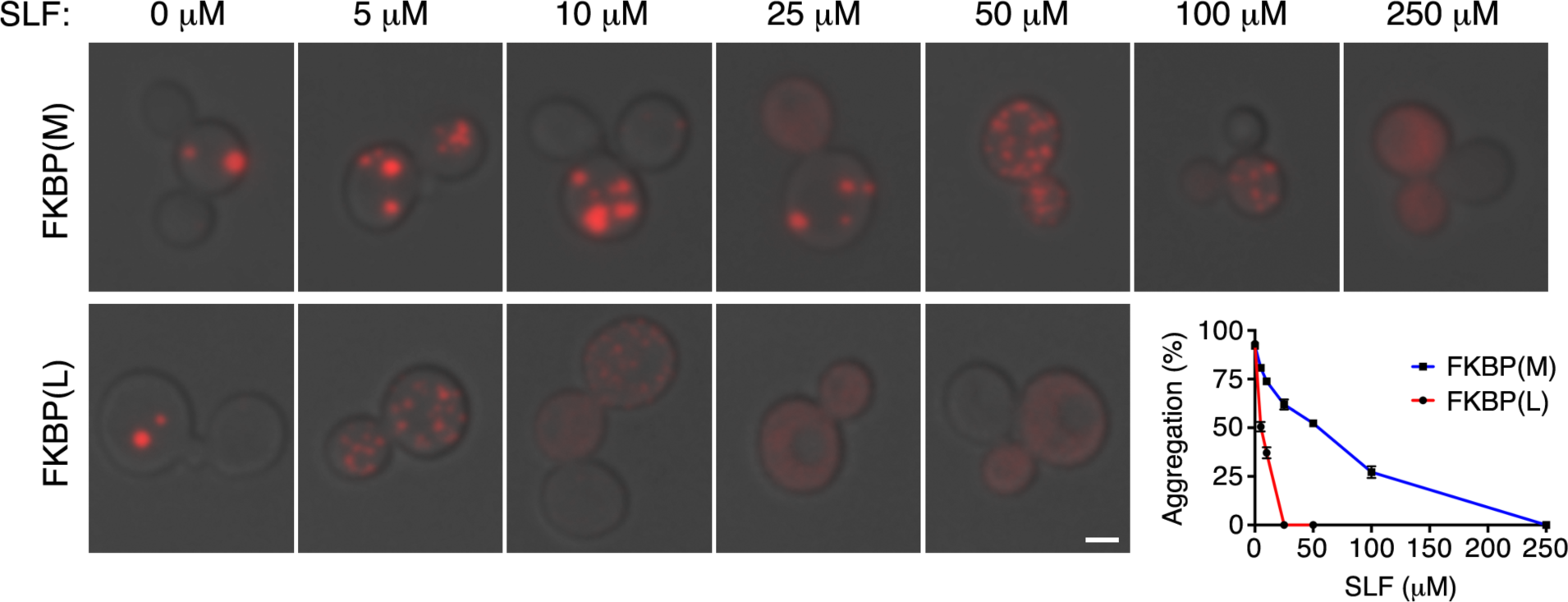
Aggregate dissolution at lower ligand concentrations with the F36L mutation. Integrating vectors encoding E2-Crimson-FKBP(M) or E2-Crimson-FKBP(L) were transformed into a *pdr1Δ pdr3Δ* strain. Logarithmically growing cultures were treated with the indicated concentrations of SLF for 30 min, and were imaged by confocal microscopy to capture red fluorescence and DIC images, using the same parameters in all cases. For each strain and condition, approximately 30 - 100 cells were examined, and the percentage of the fluorescence signal that was punctate was quantified as described in Materials and Methods. Representative merged images are shown together with quantitation of the image data. Scale bar, 2 μm. Error bars represent s.e.m.

The prediction was that E2-Crimson-FKBP(L) aggregates would dissolve at lower SLF concentrations than E2-Crimson-FKBP(M) aggregates. For this experiment, *pdr1Δ pdr3Δ* cells expressing the aggregating constructs were incubated with varying concentrations of SLF for 30 min. Confocal image stacks were then acquired, and the ratio of punctate to total fluorescence was quantified. Representative images are shown in Figure 4 together with quantitation of the image data. Maximal dissolution of E2-Crimson-FKBP(M) aggregates required 250 μM SLF, whereas maximal dissolution of E2-Crimson-FKBP(L) aggregates required only 25 μM SLF. Thus, the F36L mutation is apparently preferable to F36M for the purpose of ligand-reversible dimerization.

### Acceleration of aggregate shrinkage with the I90 V mutation

After addition of SLF, aggregates generated with E2-Crimson-FKBP(M) or E2-Crimson-FKBP(L) required up to 5 min or more for complete dissolution. Because a similar time scale is seen for cargo transport through the yeast secretory pathway (Losev *et al.*, 2006) and our ultimate goal is to produce a nearly synchronous cargo wave, faster dissolution of the aggregates would be useful.

Ligand-induced disruption of the FKBP dimer presumably occurs when the dimer spontaneously dissociates to enable ligand binding. Therefore, a mutation that increases the dissociation rate of the dimer would be expected to accelerate ligand-induced shrinkage of FKBP aggregates. To identify such a mutation, we examined the crystal structure of the FKBP(M) dimer (Rollins *et al.*, 2000). Residue Ile-90 in the dimerization interface has a hydrophobic interaction with its counterpart in the other subunit. The corresponding residue is Val in many closely related FKBP proteins (Galat, 2008), suggesting that the less hydrophobic Val should be tolerated at position 90. We introduced the I90V mutation into FKBP(L) to generate FKBP(L,V).

To visualize the shrinkage of aggregates containing E2-Crimson fused to either FKBP(L) or FKBP(L,V), we sought to perform live-cell imaging after addition of SLF. Flow chambers have been described for introducing drugs during live-cell imaging of yeast (Lee *et al.*, 2008; Charvin *et al.*, 2010), but those systems were constructed using the hydrophobic polymer polydimethylsiloxane (PDMS), which binds drugs such as SLF (Zhou *et al.*, 2012). We therefore modified and simplified a commercially available yeast live-cell imaging system (Charvin *et al.*, 2010) to avoid the use of PDMS. As shown in Figure 5A, yeast cells attached to a coverslip with concanavalin A are overlaid with a slab of the hard plastic poly(methyl methacrylate) (PMMA). A channel in the PMMA creates a flow chamber, which is connected to metal tubes that are connected in turn to plastic tubing. Liquid is pulled through the chamber by a nonelectric pressure-driven pump (Moscovici *et al.*, 2010). With this device, we can flow SLF-containing medium over the cells while performing 4D confocal microscopy.

**FIGURE 5:**
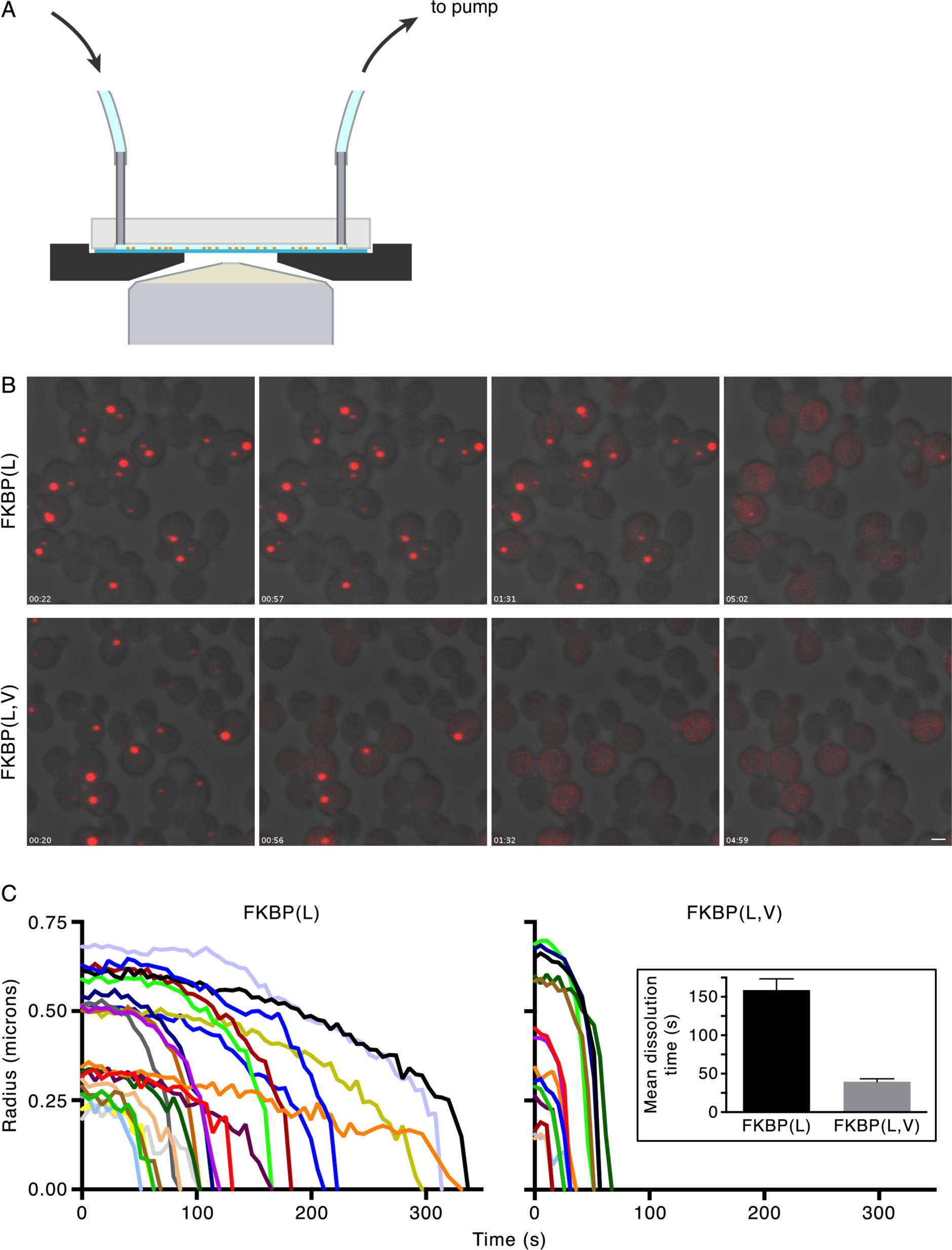
Faster aggregate dissolution with the I90V mutation. (A) Diagram of a cross-section through the center of the flow chamber. Media enters and exits the chamber through Tygon tubing attached to stainless steel tubes embedded in a slab of hard PMMA plastic. Those tubes connect to a groove in the plastic. The PMMA slab is placed on a concanavalin A-coated coverslip to which yeast cells have been attached. This assembly sits in a well in a metal piece on a microscope stage. To hold the assembly in place, an additional clear plastic slab, not shown in the diagram, is placed on top and attached with screws to the metal piece. A tapered opening in the metal piece allows an oil objective to be positioned close to the bottom of the coverslip. Medium is pulled through the flow chamber using a negative pressure pump. Hydrophobic compounds such as SLF flow directly past the cells. (B) Selected frames from Videos 1 and 2, which show SLF-induced dissolution of E2-Crimson-FKBP(L) aggregates and E2-Crimson-FKBP(L,V) aggregates, respectively. SLF reached the cells approximately 20 s after the beginning of a movie. Scale bar, 2 μm. (C) Quantitation of aggregate dissolution from Videos 1 and 2. The radius of each fluorescent aggregate was measured at each time point, beginning at the time point when SLF was estimated to reach the cells. The inset shows the mean time needed for complete dissolution of aggregates. For this quantitation, a total of 43 E2-Crimson-FKBP(L) aggregates and 36 E2-Crimson-FKBP(L,V) aggregates were examined from three movies for each construct, and the times needed for complete dissolution of the aggregates were recorded. Error bars represent s.e.m. The mean dissolution times were 159 ± 15 s for E2-Crimson-FKBP(L) and 40 ± 4 for E2-Crimson-FKBP(L,V).

This flow chamber was used to track the shrinkage of cytosolic E2-Crimson-FKBP(L) and E2-Crimson-FKBP(L,V) aggregates after addition of 25 μM SLF. The stacks of confocal images for the individual time points were projected, and the projections were assembled into movies. Figure 5B shows frames from two representative movies (Videos 1 and 2). The aggregates varied considerably in size, and accordingly, they took variable amounts of time to dissolve. On average, shrinkage of aggregates was substantially faster with E2-Crimson-FKBP(L,V) than with E2-Crimson-FKBP(L). In Figure 5C, the radii of the aggregates from the two representative movies are plotted as a function of time. The inset to Figure 5C shows that with E2-Crimson-FKBP(L,V) the mean time for complete dissolution of an aggregate was only 40 sec, approximately 4 times less than with E2-Crimson-FKBP(L). Thus, the I90V mutation substantially accelerates shrinkage of aggregates.

## Conclusions

Reversible aggregates can be generated in the secretory pathway not only with dimerizing FKBP, but also with tools such as the plant photoreceptor protein UVR8, which forms photolabile homodimers (Chen *et al.*, 2013). Another way to create a secretory cargo wave is the “retention using selective hooks” (RUSH) system, which employs biotin to release a streptavidin-binding peptide-modified reporter protein from a streptavidin-modified ER-localized hook protein (Boncompain *et al.*, 2012). Each of these methods is likely to have advantages and disadvantages for particular applications. Because dimerizing FKBP has proven to be useful and flexible, we are seeking to adapt it for generating reversible aggregates in the yeast secretory pathway.

As the first step in this project, we used rational mutagenesis to improve the properties of dimerizing FKBP. The widely used FKBP(M) mutant has the F36M mutation, which promotes dimerization but also reduces ligand binding affinities (Rollins *et al.*, 2000). By contrast, the F36L mutation was reported to have only a minimal effect on ligand binding affinities (DeCenzo *et al.*, 1996), and we found that the FKBP(L) mutant dimerizes. Aggregates generated with FKBP(L) dissolve at about 10-fold lower ligand concentrations than aggregates generated with FKBP(M).

Another opportunity for improvement concerned the rate of ligand-induced disruption of the dimer. The dimer dissociates spontaneously at some rate, enabling ligand to bind and to occlude the dimerization interface. Based on this reasoning, we examined the crystal structure of the FKBP(M) dimer, and noted that hydrophobic interactions involving Ile-90 probably contribute substantially to dimerization (Rollins *et al.*, 2000). We therefore replaced Ile-90 with Val, which is structurally similar but less hydrophobic. The I90V mutation is expected to accelerate spontaneous dissociation of the dimer. Consistent with this prediction, aggregates generated with the FKBP(L,V) mutant, which has both the F36L and I90V mutations, shrink about 4-fold faster after ligand addition than aggregates generated with FKBP(L).

We have shown that fluorescent aggregates can be generated in the yeast cytosol by fusing FKBP(L,V) to the tetrameric fluorescent protein E2-Crimson. In a yeast strain with reduced expression of pleiotropic drug transporters, E2-Crimson-FKBP(L,V) aggregates can be dissolved rapidly by adding the ligand SLF. Further engineering will be required to use this sytem to create a secretory cargo wave. For example, yeast esterases can apparently degrade SLF (J. Casler and E. McLean, unpublished observations), so inhibition or removal of esterases should help to prevent reaggregation. Another concern is that because both E2-Crimson and FKBP(L,V) oligomerize, the E2-Crimson-FKBP(L,V) fusion protein is unlikely to cross the ER membrane posttranslationally, and will need to be fused to a signal sequence that drives efficient cotranslational translocation (Ng *et al.*, 1996; Fitzgerald and Glick, 2014). We are working to overcome these challenges in order to engineer a regulatable fluorescent secretory cargo for yeast cells.

## Materials and Methods

### Strains, plasmids, and reagents

The parental haploid *S. cerevisiae* strain was JK9-3da (*leu2-3,112 ura3-52 rme1 trp1 his4*) (Kunz *et al.*, 1993). Yeast cells were grown in minimal glucose dropout medium (SD) (Sherman, 1991) or nonfluorescent minimal medium (NSD) (Bevis *et al.*, 2002) prepared using nutrient mixtures from Sunrise Science Products (San Diego, CA). The *PDR5* and *PDR1* genes were disrupted by PCR-based gene deletion using a kanamycin resistance cassette (Longtine *et al.*, 1998), and the *PDR3* gene was disrupted in a similar manner using a nourseothricin resistance cassette (Goldstein and McCusker, 1999).

DNA constructs were designed with the aid of SnapGene software (GSL Biotech, Chicago, IL), and the Supplemental Materials includes a folder of five annotated plasmid map/sequence files that can be opened with SnapGene Viewer (www.snapgene.com/products/snapgeneviewer). Expression with the integrating vector YIplac204 (Gietz and Sugino, 1988) was driven by the strong constitutive *TPI1* promoter and the *CYC1* terminator (Losev *et al.*, 2006). Plasmids were modified by standard methods including primer-directed mutagenesis and Gibson assembly, and critical regions were verified by sequencing. Integrating vectors were linearized and then transformed into yeast cells using lithium acetate (Gietz and Woods, 2002) for integration at the *TRP1* locus. Single-copy integrations were confirmed by PCR using primers 5’-GTGTACTTTGCAGTTATGACGCCAGATGG-3’ and 5’-AGTCAACCCCCTGCGATGTATATTTTCCTG-3’.

A 100 mM solution of SLF in ethanol was obtained from Cayman Chemical (Ann Arbor, MI). This drug was diluted in NSD to the desired concentration.

### Construction of the flow chamber

A Warner Instruments model YC-1 yeast cell flow chamber, which was based on a published design (Charvin *et al.*, 2010), was modified and simplified as follows. The device was supplied with a PDMS flow chamber containing 1.27-mm outer diameter stainless steel perfusion port tubes that terminated in a 100-μm deep groove in the chamber. Those tubes were removed, and were inserted into a custom fabricated PMMA slab of the same size with a similar groove. The YC-1 chamber immobilizes yeast cells by gently compressing them between a cellulose membrane and a PDMS-coated coverslip. Instead, we omitted the cellulose membrane, and immobilized the cells on a 24x50 mm high precision No. 1.5 coverslip (Marienfeld, Lauda-Königshofen, Germany) that had been coated with concanavalin A as described below under “Fluorescence microscopy”. The assembly was sealed with a clear top plate that was screwed into place. With this configuration, the medium flowed directly over the cells and never encountered PDMS.

Medium was flowed into the chamber through 1/32” Tygon tubing connected to the stainless steel perfusion port tubes. To minimize leakage, the flow was driven using negative pressure. We employed an inexpensive nonelectric pressure-driven pump (Moscovici *et al.*, 2010), with a tank volume created using four 140-cc syringes (Healthcare Supply Pros, Austin, TX) that were clamped with the plungers in an extended position using 1.5 x 4 inch EZ White Snap Clamps (Amazon.com). Pressure was created using a 60-cc syringe that was clamped with the plunger in an extended position using a 0.75 x 4 inch EZ White Snap Clamp. This setup allowed medium to be flowed over the cells for 10-15 minutes at a rate of approximately 1 mL/min. Presumably, similar results could be obtained using a more conventional electric syringe pump to generate negative pressure.

### Fluorescence microscopy

Static images of cells were captured as Z-stacks using a Zeiss LSM 880 confocal microscope equipped with a 1.4-NA 63x oil objective. The Z-stacks were average projected using ImageJ (https://imagej.nih.gov/ij/).

For 4D live-cell imaging (Day *et al.*, 2016), a 5-mL yeast culture was grown overnight in NSD in a baffled flask, and was diluted to an OD_600_ of 0.5 - 0.7 in fresh NSD medium. A 24x50 mm coverslip was coated with freshly dissolved 2 mg/mL concanavalin A (Sigma-Aldrich, St. Louis, MO; product number C2010) for 10 min, then washed with water and allowed to dry. A 250-μL aliquot of the culture was overlaid on the coverslip, and cells were allowed to adhere for 10 min, followed by several rounds of gentle washing with NSD. The coverslip was placed in the flow chamber and sealed while ensuring that the cells remained immersed in NSD the entire time. NSD was pushed through the flow chamber with a syringe to remove any bubbles, and then the Tygon tubing was clamped. The flow chamber was placed on the stage of a Leica SP5 inverted confocal microscope equipped with a 1.4-NA 63x oil objective. One end of the Tygon tubing was connected to the negative pressure pump while the other end was placed in a culture tube with SLF-containing medium. To initiate flow of medium, the clamp was removed. Z-stacks were captured at intervals of 5-6 sec.

Individual images were resized, cropped, and processed to adjust brightness and contrast using Adobe Photoshop. To quantify the fraction of a fluorescent construct that was aggregated at a given SLF concentration, a custom ImageJ plugin was used as described in the Supplemental Materials. 4D data sets were processed into videos using ImageJ (Day *et al.*, 2016). To quantify the radius of a fluorescent aggregate at a given time point in a video, a maximum intensity projection of the confocal image stack was processed with ImageJ by calibrating the image in microns, setting a threshold that excluded cytoplasmic signal, and using the Analyze Particles tool to measure the area of the aggregate. The radius was then calculated by dividing the square root of the area by π. To determine the time needed for dissolution of a fluorescent aggregate, the aggregate was tracked using maximum intensity projections until it was no longer visible.

## Abbreviations

DIC, differential interference contrast; EGFP, enhanced GFP; FKBP, FK506-binding protein; NSD, nonfluorescent synthetic glucose medium; PDMS, polydimethylsiloxane; PMMA, poly(methyl methacrylate); SLF, synthetic ligand of FKBP

## Acknowledgments

This work was supported by NIH grant R01 GM104010. Development of the flow chamber was supported by funding through the Biological Systems Science Division, Office of Biological and Environmental Research, Office of Science, U.S. Dept. of Energy, under Contract DE-AC02-06CH11357. The negative pressure pump was constructed with the assistance of Cari Launiere at the Microenvironmental Control Foundry at the University of Illinois, Chicago, a facility sponsored by the Chicago Biomedical Consortium with support from the Searle Funds at The Chicago Community Trust. JCC and KJD were supported by NIH training grant T32 GM007183. Fluorescence microscopy was performed with the assistance of Vytas Bindokas at the Integrated Microscopy Core Facility, which received support from the NIH-funded Cancer Center Support Grant P30 CA014599. Fernando Valbuena helped to optimize the PCR-based method for confirming single-copy integrations.

## Supplemental Videos

**Video 1.**

Dissolution of E2-Crimson-FKBP(L) aggregates after SLF-containing medium was flowed over the cells. Confocal Z-stacks were collected at intervals of 5.162 s. This movie was used to generate data shown in the upper panels of Figure 5B and the left panel of Figure 5C.

**Video 2.**

Dissolution of E2-Crimson-FKBP(L,V) aggregates after SLF-containing medium was flowed over the cells. Confocal Z-stacks were collected at intervals of 5.705 s. This movie was used to generate data shown in the lower panels of Figure 5B and the right panel of Figure 5C.

## References

Bevis, B.J., Hammond, A.T., Reinke, C.A., and Glick, B.S. (2002). *De novo* formation of transitional ER sites and Golgi structures in *Pichia pastoris*. Nat. Cell Biol. 4, 750–756.

Boncompain, G., Divoux, S., Gareil, N., de Forges, H., Lescure, A., Latreche, L., Mercanti, V., Jollivet, F., Raposo, G., and Perez, F. (2012). Synchronization of secretory protein traffic in populations of cells. Nat. Methods 9, 493–498.

Charvin, G., Oikonomou, C., and Cross, F. (2010). Long-term imaging in microfluidic devices. Methods Mol. Biol. 591, 229–242.

Chen, D., Gibson, E.S., and Kennedy, M.J. (2013). A light-triggered protein secretion system. J. Cell Biol. 201, 631–640.

Day, K.J., Papanikou, E., and Glick, B.S. (2016). 4D confocal imaging of yeast organelles. Methods Mol. Biol., in press.

DeCenzo, M.T., Park, S.T., Jarrett, B.P., Aldape, R.A., Futer, O., Murcko, M.A., and Livingston, D.J. (1996). FK506-binding protein mutational analysis: defining the active-site residue contributions to catalysis and the stability of ligand complexes. Protein Eng. 9, 173–180.

Fegan, A., White, B., Carlson, J.C., and Wagner, C.R. (2010). Chemically controlled protein assembly: techniques and applications. Chem. Rev. 110, 3315–3336.

Fitzgerald, I., and Glick, B.S. (2014). Secretion of a foreign protein from budding yeasts is enhanced by cotranslational translocation and by suppression of vacuolar targeting. Microb. Cell Fact. 13.

Galat, A. (2008). Functional drift of sequence attributes in the FK506-binding proteins (FKBPs). J. Chem. Inf. Model. 48, 1118–1130.

Galat, A. (2013). Functional diversity and pharmacological profiles of the FKBPs and their complexes with small natural ligands. Cell. Mol. Life Sci. 70, 3243–3275.

Gietz, R.D., and Sugino, A. (1988). New *yeast-Escherichia coli* shuttle vectors constructed with in vitro mutagenized yeast genes lacking six-base pair restriction sites. Gene 74, 527–534.

Gietz, R.D., and Woods, R.A. (2002). Transformation of yeast by lithium acetate/single-stranded carrier DNA/polyethylene glycol method. Methods Enzymol. 350, 87–96.

Goldstein, A.L., and McCusker, J.H. (1999). Three new dominant drug resistance cassettes for gene disruption in *Saccharomyces cerevisiae*. Yeast 15, 1541–1553.

Holt, D.A., Luengo, J.I., Yamashita, D.S., Oh, H.J., Konialian, A.L., Yen, H.K., Rozamus, L.W., Brandt, M., Bossard, M.J., Levy, M.A., Eggleston, D.S., Liang, J., Schultz, L.W., Stout, T.J., and Clardy, J. (1993). Design, synthesis, and kinetic evaluation of high-affinity FKBP ligands and the X-ray crystal structures of their complexes with FKBP12. J. Am. Chem. Soc. 115, 9925–9938.

Kaiser, C.A., Gimeno, R.E., and Shaywitz, D.A. (1997). Protein secretion, membrane biogenesis, and endocytosis. In: The Molecular and Cellular Biology of the Yeast Saccharomyces, vol. 3, eds. J.R. Pringle, J.R. Broach, and E.W. Jones: Cold Spring Harbor Laboratory Press, 91–227.

Kunz, J., Schneider, U., Deuter-Reinhard, M., Movva, N.R., and Hall, M.N. (1993). Target of rapamycin in yeast, TOR2, is an essential phosphatidylinositol kinase homolog required for G1 progression. Cell 73, 585–596.

Lavieu, G., Zheng, H., and Rothman, J.E. (2013). Stapled Golgi cisternae remain in place as cargo passes through the stack. eLife 2, e00558.

Lee, P.J., Helman, N., C., Lim, W.A., and Hung, P.J. (2008). A microfluidic system for dynamic yeast cell imaging. BioTechniques 44, 91–95.

Lisenbee, C.S., Karnik, S.K., and Trelease, R.N. (2003). Overexpression and mislocalization of a tail-anchored GFP redefines the identity of peroxisomal ER. Traffic 4, 491–501.

Longtine, M.S., McKenzie, A., 3rd, Demarini, D.J., Shah, N.G., Wach, A., Brachat, A., Philippsen, P., and Pringle, J.R. (1998). Additional modules for versatile and economical PCR-based gene deletion and modification in *Saccharomyces cerevisiae*. Yeast 14, 953–961.

Losev, E., Reinke, C.A., Jellen, J., Strongin, D.E., Bevis, B.J., and Glick, B.S. (2006). Golgi maturation visualized in living yeast. Nature 22, 1002–1006.

Moscovici, M., Chien, W.Y., Abdelgawad, M., and Sun, Y. (2010). Electrical power free, low dead volume, pressure-driven pumping for microfluidic applications. Biomicrofluidics 4, 46501.

Ng, D.T., Brown, J.D., and Walter, P. (1996). Signal sequences specify the targeting route to the endoplasmic reticulum membrane. J. Cell Biol. 134, 269–278.

Papanikou, E., and Glick, B.S. (2009). The yeast Golgi apparatus: insights and mysteries. FEBS Lett. 583, 3746–3751.

Rivera, V.M., Wang, X., Wardwell, S., Courage, N.L., Volchuk, A., Keenan, T., Holt, D.A., Gilman, M., Orci, L., Cerasoli, F.J., Rothman, J.E., and Clackson, T. (2000). Regulation of protein secretion through controlled aggregation in the endoplasmic reticulum. Science 287, 826–830.

Rizzo, R., Parashuraman, S., Mirabelli, P., Puri, C., Lucocq, J., and Luini, A. (2013). The dynamics of engineered resident proteins in the mammalian Golgi complex relies on cisternal maturation. J. Cell Biol. 201, 1027–1036.

Rogers, B., Decottignies, A., Kolaczkowski, M., Carvajal, E., Balzi, E., and Goffeau, A. (2001). The pleitropic drug ABC transporters from *Saccharomyces cerevisiae*. J. Mol. Microbiol. Biotechnol. 3, 207–214.

Rollins, C.T., Rivera, V.M., Woolfson, D.N., Keenan, T., Hatada, M., Adams, S.E., Andrade, L.J., Yaeger, D., van Schravendijk, M.R., Holt, D.A., Gilman, M., and Clackson, T. (2000). A ligand-reversible dimerization system for controlling protein-protein interactions. Proc. Natl. Acad. Sci. USA 97, 7096–7101.

Schüller, C., Mamnun, Y.M., Wolfger, H., Rockwell, N., Thorner, J., and Kuchler, K. (2007). Membrane-active compounds activate the transcription factors Pdr1 and Pdr3 connecting pleiotropic drug resistance and membrane lipid homeostasis in*Saccharomyces cerevisiae*. Mol. Biol. Cell 18, 4932–4944.

Sherman, F. (1991). Getting started with yeast. Methods Enzymol. 194, 3–21.

Snapp, E.L., Hegde, R.S., Francolini, M., Lombardo, F., Colombo, S., Pedrazzini, E., Borgese, N., and Lippincott-Schwartz, J. (2003). Formation of stacked ER cisternae by low affinity protein interactions. J. Cell Biol. 163, 257–269.

Strack, R.L., Hein, B., Bhattacharyya, D., Hell, S.W., Keenan, R.J., and Glick, B.S. (2009).A rapidly maturing far-red derivative of DsRed-Express2 for whole-cell labeling. aBiochemistry 48, 8279–8281.

Thakur, J.K., Arthanari, H., Yang, F., Pan, S.J., Fan, X., Breger, J., Frueh, D.P., Gulshan, K., Li, D.K., Mylonakis, E., Struhl, K., Moye-Rowley, W.S., Cormack, B.P., Wagner, G., and Näär, A.M. (2008). A nuclear receptor-like pathway regulating multidrug resistance in fungi. Nature 452, 604–609.

Volchuk, A., Amherdt, M., Ravazzola, M., Brugger, B., Rivera, V.M., Clackson, T., Perrelet, A., Söllner, T.H., Rothman, J.E., and Orci, L. (2000). Megavesicles implicated in the rapid transport of intracisternal aggregates across the Golgi stack. Cell 102, 335–348.

Zacharias, D.A., Violin, J.D., Newton, A.C., and Tsien, R.Y. (2002). Partitioning of lipid-modified monomeric GFPs into membrane microdomains of live cells. Science 296, 913–916.

Zhou, J., Khodakov, D.A., Ellis, A.V., and Voelcker, N.H. (2012). Surface modification for PDMS-based microfluidic devices. Electrophoresis 33, 89–104.

